# Fast Kernel-based Association Testing of non-linear genetic effects for Biobank-scale data

**DOI:** 10.1101/2022.04.13.488214

**Authors:** Boyang Fu, Ali Pazokitoroudi, Mukund Sudarshan, Lakshminarayanan Subramanian, Sriram Sankararaman

**Affiliations:** Department of Computer Science, UCLA, Los Angeles, CA, USA; Department of Human Genetics, David Geffen School of Medicine, UCLA, Los Angeles, CA, USA; Department of Computational Medicine, David Geffen School of Medicine, UCLA, Los Angeles, CA, USA; Department of Computer Science, New York University, New York, NY, USA; Department of Population Health, NYU School of Medicine, New York, NY, USA

## Abstract

Our knowledge of non-linear genetic effects on complex traits remains limited, in part, due to the modest power to detect such effects. While kernel-based tests offer a powerful approach to test for nonlinear relationships between sets of genetic variants and traits, current approaches cannot be applied to Biobank-scale datasets containing hundreds of thousands of individuals. We propose, FastKAST, a Kernel-based approach that can test for non-linear effects of a set of variants on a trait. FastKAST provides calibrated hypothesis tests while enabling analysis of Biobank-scale datasets with hundreds of thousands of individuals. We applied FastKAST to thirty quantitative traits measured across ≈ 300 K unrelated white British individuals in the UK Biobank to detect sets of variants with nonlinear effects at genome-wide significance.

## Introduction

Understanding the contribution of non-linear genetic effects on complex traits is a central question in human genetics [1–7]. A powerful approach to identify such effects relies on grouping genetic variants into “sets” and testing their aggregated effect [8–13]. The mixed model framework offers a versatile approach to test such effects: capable of testing a wide range of linear and non-linear relationships between genotype and trait by employing a kernel function that measures similarity between pairs of genotypes [11–13]. In practice, testing within the mixed model framework is computationally impractical for large datasets so that current approaches typically restrict their focus to linear additive models [11]. While biobank-scale datasets containing genetic and phenotypic data over hundreds of thousands of individuals provide the large sample sizes needed to identify non-linear effects [14–16], computational challenges have limited these efforts.

We propose Fast non-linear Kernel-based ASsociation Test (FastKAST), a scalable approach to test for non-linear effects of a set of variants on a trait in a mixed model framework. Specifically, FastKAST permits fitting of a wide-class of kernel functions that model non-linear effects of genetic variants on a trait (including the popular radial basis function (RBF) kernel). FastKAST combines a low-dimensional approximation to the kernel function [17] within a score test, obtaining calibrated p-values by fitting a distribution to genomewide statistics obtained from a small number of permuted phenotypes [18]. As a result, FastKAST can efficiently test nonlinear associations in biobank-scale data.

Our theoretical and empirical analyses show that FastKAST provides calibrated hypothesis tests. Using extensive simulations across genetic architectures in which the phenotypes have a linear dependence on genotype (consistent with the known polygenic architecture of most complex phenotypes [19–21]) but no non-linear dependencies, we find that FastKAST provides calibrated p-values. On small-scale datasets that permit exact computation, FastKAST is highly concordant with exact tests. To illustrate its utility, we applied FastKAST with a RBF kernel to thirty quantitative traits measured across *N* ≈ 300K unrelated white British individuals in the UK Biobank (UKBB). Testing genotypes measured on the UKBB array across 100kb windows, we find 27 windows with statistically significant nonlinear effects in seven traits (*p* < 6 × 10^−8^ accounting for the number of sets and traits tested). To further interrogate the nature of these effects, we repeated our analyses after regressing out pairwise interactions (in addition to linear effects) and on imputed genotypes in the UKBB to find two significant associations for blood Mean Platelet Volume (MPV) and serum urate levels. Our results highlight the potential of FastKAST to uncover non-linear genetic effects from Biobank-scale datasets.

## Results

### Methods overview

FastKAST tests for non-linear effects between genotypes measures on a set of *M* single nucleotide polymorphisms (SNPs) and a phenotype. The vector of phenotypes ***y***, measured on *N* individuals, is modeled as:

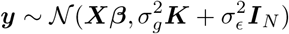

Here ***X*** denotes fixed effects. ***K*** is a *N* × *N* kernel matrix obtained by applying a kernel function *k* to every pair of genotypes over the *M* SNPs to be tested, *i*.*e*., entry (*i, j*) in the matrix ***K***, *K*_*i,j*_ = *k*(***z***_*i*_, ***z***_*j*_) where ***z***_*i*_ (***z***_*j*_) denotes the genotype of individual *i* (*j*). 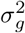 denotes the variance component associated with genetic effects while 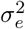 denotes the variance component associated with residual effects. The kernel function can model different relationships between genotype and phenotype: the inner-product kernel 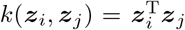 implies a linear additive model while the radial basis function (RBF) kernel 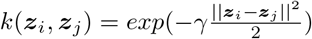 is a common kernel to model non-linear relationships.

Testing for a genetic contribution in this model involves testing the null hypothesis: 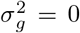 which is commonly achieved using the score test [11]. While p-values for the score test can be computed efficiently when testing linear effects (as implemented in the SKAT software [11]), these approaches are computationally impractical for testing non-linear effects in large samples.

FastKAST approximates the kernel function by transforming the input genotypes to a randomized feature space [17] where the number of random features *D* (termed the approximation dimension) determines the quality of the approximation. Combining the idea that an approximation dimension *D* substantially smaller than the sample size *N* is sufficient for approximating the kernel function with efficient linear algebra implementations allows FastKAST to efficiently compute p-values. While these p-value computations assume that the kernel hyperparameters are known (*e*.*g*., the *γ* parameter for the RBF kernel), the more common setting is one in which the hyperparameter is unknown. In this more general setting (which is the setting that we focus on in this work), FastKAST adaptively selects the hyperparameter and obtains calibrated p-values by fitting a distribution to genome-wide statistics obtained from a small number of permuted phenotypes [18] (see Methods for details).

### Calibration of FastKAST

To assess the calibration of FastKAST, we performed simulations of quantitative traits with linear additive effects but no non-linear effects. We simulated phenotypes based on whole-genome genotypes from unrelated white British individuals in the UK Biobank (UKBB) (*N* = 337, 205 individuals and *M* = 593, 300 SNPs on the UK Biobank Axiom array; see Methods for details on dataset). We performed simulations under four genetic architectures: infinitesimal model (causal variants ratio = 1); non-infinitesimal model (causal variants ratio = 0.001) with a different range of minor allele frequencies (MAF) for the causal variants: [0.01, 0.05] (RARE), [0.05-0.5] (COMMON), [0, 0.5] (ALL). The trait heritability was set to *h*^2^ = 0.50 in all settings.

We applied FastKAST with the radial basis function (RBF) kernel in non-overlapping 100 kb windows. We approximated the RBF kernel with approximation dimension *D* = 50*M* where *M* is the number of SNPs within each window. Since the goal of our work is to identify sets of SNPs with non-linear effects, we need to first completely regress out the linear effect before testing for non-linear effects. We observe that simply regressing out the effects of SNPs in the set being tested does not yield calibrated tests likely due to correlation or linkage disequilibrium (LD) across SNPs (Figure S1). On the other hand, regressing out the linear effect within the target window as well as the additional neighboring windows can solve this issue (termed superwindow, the size of which is measured in multiples of the target window size). We empirically show that a superwindow of size five (*e*.*g*., target window plus two neighboring windows on each side) leads to calibrated p-values and appropriate control of the false positive rate (Figure S1). With this approach to control for linear effects, FastKAST obtains calibrated p-values across the architectures considered (Figure 1(a); Table S1). While FastKAST adaptively chooses the kernel hyperparameter, our theory (Supplementary Note S1) and empirical results show that FastKAST remains calibrated even for a specific choice of hyperparameter (Figure S2, S3; Table S2).

**Figure 1:**
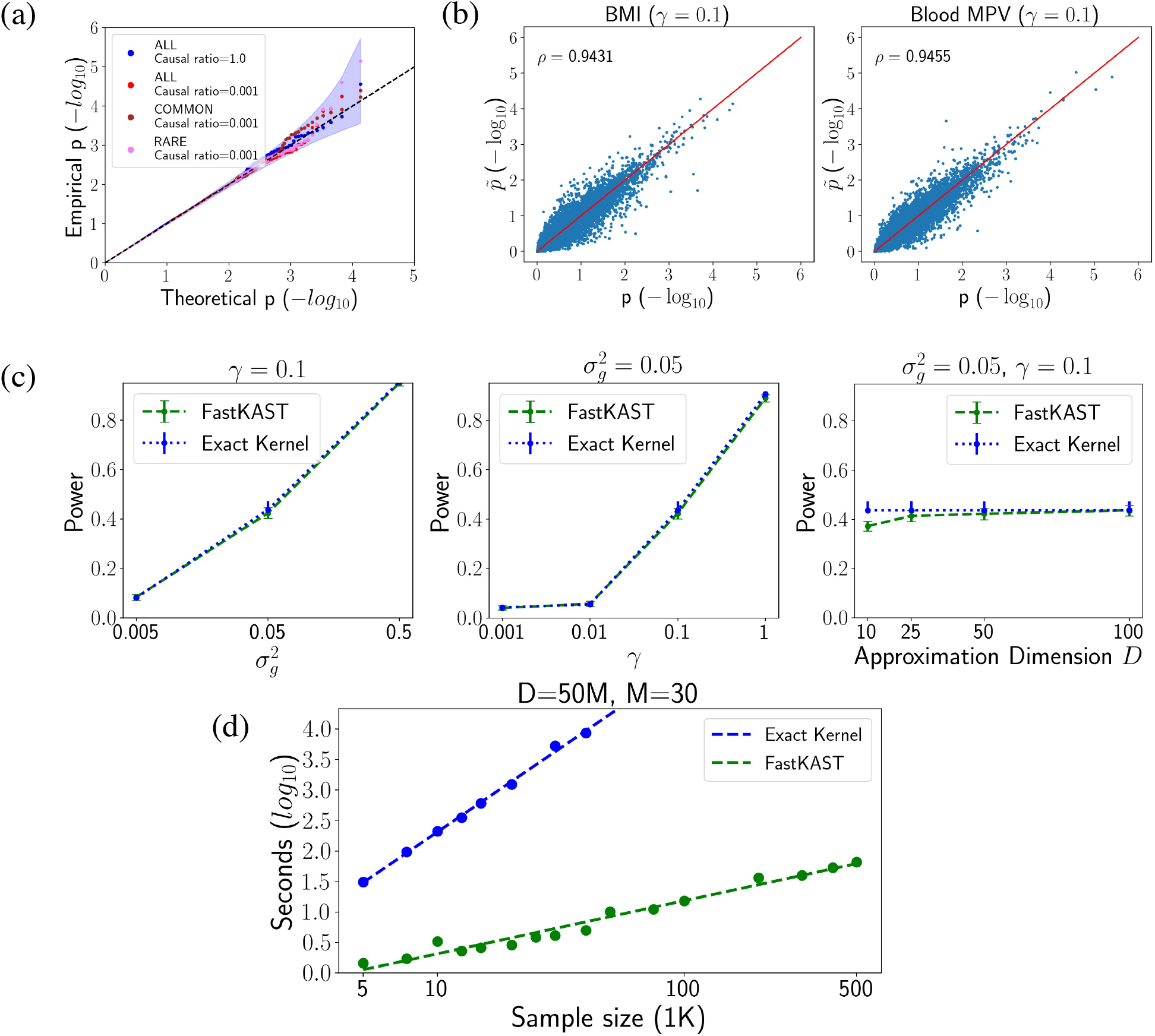
Assessment of calibration, power and scalability of FastKAST. **(a)** Calibration of FastKAST under null simulations that include linear effects but no non-linear effects (*N* = 50*K* individuals). We fixed SNP heritability at 0.50 while varying the ratio of causal variants (∈ {0.001, 1}) and the range of minor allele frequencies (MAF) of the causal variants (ALL, COMMON, RARE). We applied FastKAST to test for nonlinear effects within 100 kb windows (after regressing out the linear effect in five windows centered around the tested window). **(b)** Comparison of p-values computed using FastKAST to an exact method. We analyzed BMI and blood Mean Platelet Volume (MPV) across 5, 000 unrelated white British individuals in the UK Biobank (UKBB). We tested each trait for non-linear effects of SNPs in the UKBB genotyping array within non-overlapping 100 kb windows using the exact RBF kernel and FastKAST (with approximation dimension *D* = 50*M* where *M* is the number of SNPs in a tested window and the kernel hyperparameter *γ* = 0.1). We regressed out linear effects among SNPs in five windows centered around the tested window while also regressing out the fixed effects associated with sex, age, and the first 20 genetic PCs. **(c)** Power of FastKAST as a function of kernel hyperparameter *γ*, kernel variance component 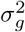, and the approximation dimension *D*. **(d)** Run time of FastKAST and the exact method as a function of sample size (*N*). The exact method requires hours to analyze sample sizes larger than 50K. FastKAST remains efficient for sample sizes as large as 500K.

### Power analysis of FastKAST

Our next experiment sought to compare the p-values obtained by FastKAST to an exact test. In the first set of experiments, we analyzed the correlation in p-values between an exact test using the RBF kernel with hyperparameter *γ* = 0.1 and the approximate kernel used by FastKAST in a simulation with causal variant ratio = 0.001 with *h*^2^ = 0.5. We limited our sample size to 8000 individuals due to limitations of computing the exact kernel. Since the approximation accuracy depends on the approximation dimension *D*, we explored the correlation between exact test p-values and FastKAST p-values by varying *D*. We found that values of *D* ≥ 50*M*, where *M* is the number of SNPs in the set, resulted in consistently high correlation (≥ 0.9) (Fig S4). To further validate our choice of the approximation dimension, we compared p-values from a test employing the exact kernel (RBF kernel with hyperparameter *γ* = 0.1) with FastKAST (*D* = 50*M*) on two real traits: Body mass index (BMI) and blood MPV. Applying both tests to assess non-linear effects within 100kb windows across 5000 unrelated white British individuals, p-values obtained by FastKAST are highly correlated with those obtained by the exact test (Pearson correlation *ρ* of 0.94 for both traits; Figure 1b). These results remain consistent across values of the RBF kernel hyperparameter *γ* and for different runs of FastKAST (Figures S5, S6).

We compared the statistical power of FastKAST relative to an exact test using simulated phenotypes with true non-linear effects. We varied the RBF kernel hyperparameter *γ* (a measure of the scale of nonlinearity), the kernel variance component 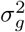 (a measure of the strength of the non-linear signal), and the approximation dimension *D* (with default values of *D* = 50M, *γ* = 0.1, and 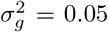). For each setting, we randomly selected 2000 windows of length 100 kb across 5, 000 individuals and simulated phenotypes 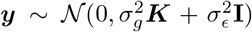, where ***K*** is constructed using the RBF kernel with hyperparameter *γ*. Across these simulations, the power of FastKAST is indistinguishable from that of the exact test provided the approximation dimension *D* ≥ 50*M* (Figure 1c). Based on these results, we decided to use *D* = 50*M* as our approximation dimension across the remaining experiments.

### Computational efficiency of FastKAST

We compared the runtime of FastKAST to the exact test with increasing sample size with the number of SNPs set to *M* = 30 (about the average number of SNPS in a 100 kb when analyzing SNPs from the UKBB genotyping array) and *D* = 50*M*. For each setting (a given set of *N, D*), we randomly subsampled *N* individuals from UKBB and *M* consecutive SNPs and reported the average runtime across 100 runs (10 replicates for sample sizes larger than 30K).

The exact test has a runtime that increases rapidly with sample size: requiring more than five hours to analyze *N* = 50*K* (Figure 1d) and extrapolated to require over 100 days to analyze *N* = 500*K* samples. On the other hand, even on the largest sample size of *N* = 500*K* (with *M* = 30,*D* = 50*M*), FastKAST requires less than five minutes to analyze a single set (this includes the time to compute p-values across multiple hyperparameter values and to analyze permuted phenotypes). We also found that the runtime of FastKAST scales quadratically as a function of the number of SNPs and approximation dimension so that FastKAST is best suited for analyzing relative small sets of SNPs (Figure S7)

### Application of FastKAST to identify non-linear effects in the UK Biobank

We applied FastKAST to about 300K unrelated white British individuals in the UKBB. We tested nonoverlapping 100 kb windows (considering SNPs with MAF > 1% in the UKBB genotyping array) to test for non-linear effects using the RBF kernel and each of thirty quantitative traits (see Methods for details on data processing). For each window tested, we regressed out linear effect of genotypes using a superwindow of size five. We included sex, age, and the top 20 genetic principal components (PCs) as covariates in all our analyses. We screened potentially interesting phenotypes using a fixed hyperparameter *γ* = 0.1 and selected traits with at least one p-value that passed the significance threshold (*p* < 6 × 10^−8^ accounting for the number of sets and traits tested). For the seven traits that passed this screening criterion, we applied FastKAST with an adaptive search for hyperparameter to obtain a p-value for each window.

We detected 27 statistically significant associations (*p* < 6 × 10^−8^) for seven traits: serum urate levels, sex hormone-binding globulin (SHBG), cystatin C, blood mean sphered cell volume (blood MSCV), blood mean platelet volume (blood MPV), blood mean corpuscular hemoglobin (blood MCH), and Low-density lipoprotein (LDL) (Figure 2, Figure S8, Figure S9). We tested the stability of our findings by recomputing p-values using FastKAST across the significant loci to find that 26 out of 27 loci remain significant (locus 22.8 that we found originally to be associated with cystatin C obtains a p-value of 1.2 × 10^−5^). We further assessed the robustness of our results to population structure by varying the number of PCs included (from five to forty) and found the statistical significance to be stable to the choice of number of PCs (Table S3).

**Figure 2:**
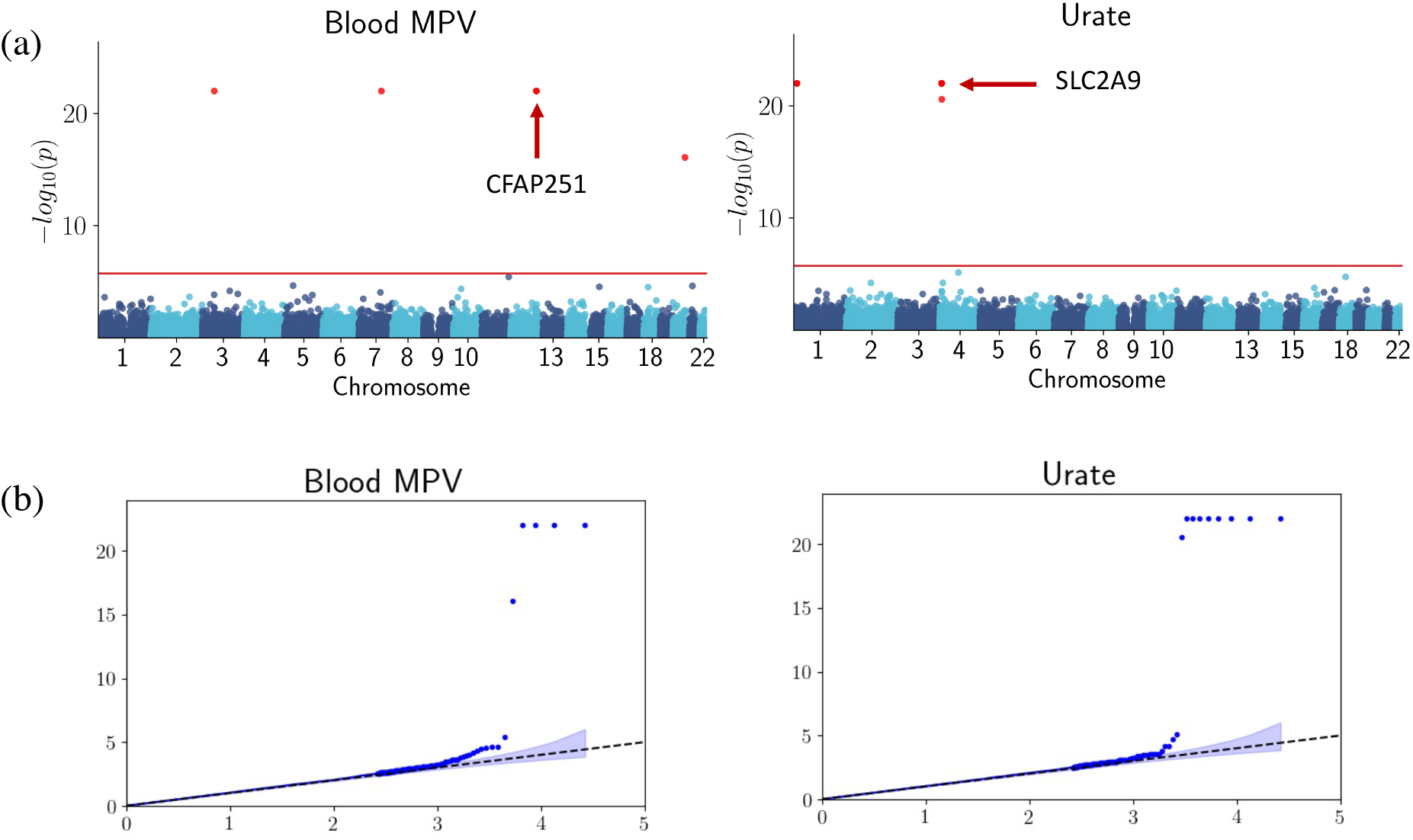
Traits that demonstrate strong evidence for nonlinear genetic effects. (a) Manhattan plot for test of non-linear effects of Blood MPV and Urate in the UK Biobank. (b). QQ-plot of the corresponding traits.

We performed additional analyses to investigate the nature of the non-linear effects at these loci. First, we repeated our analysis by regressing out linear and quadratic effects and repeated the test using FastKAST (“Non-linear+non-quadratic” column in Table 1). Previous studies have shown that apparent non-linear genetic effects could potentially be explained by a model of linear effects involving untyped causal variants and correlation between tested genotypes with untyped causal variants [22]. We investigated this possibility by testing the significant loci using imputed genotypes (column “Non-linear (imputed)” in Table 1). We found that 4 out of the 27 loci remain significant using imputed genotypes of which two loci (one associated with blood MPV and the other with serum urate levels) remain significant after removing both linear and quadratic effects.

**Table 1:**
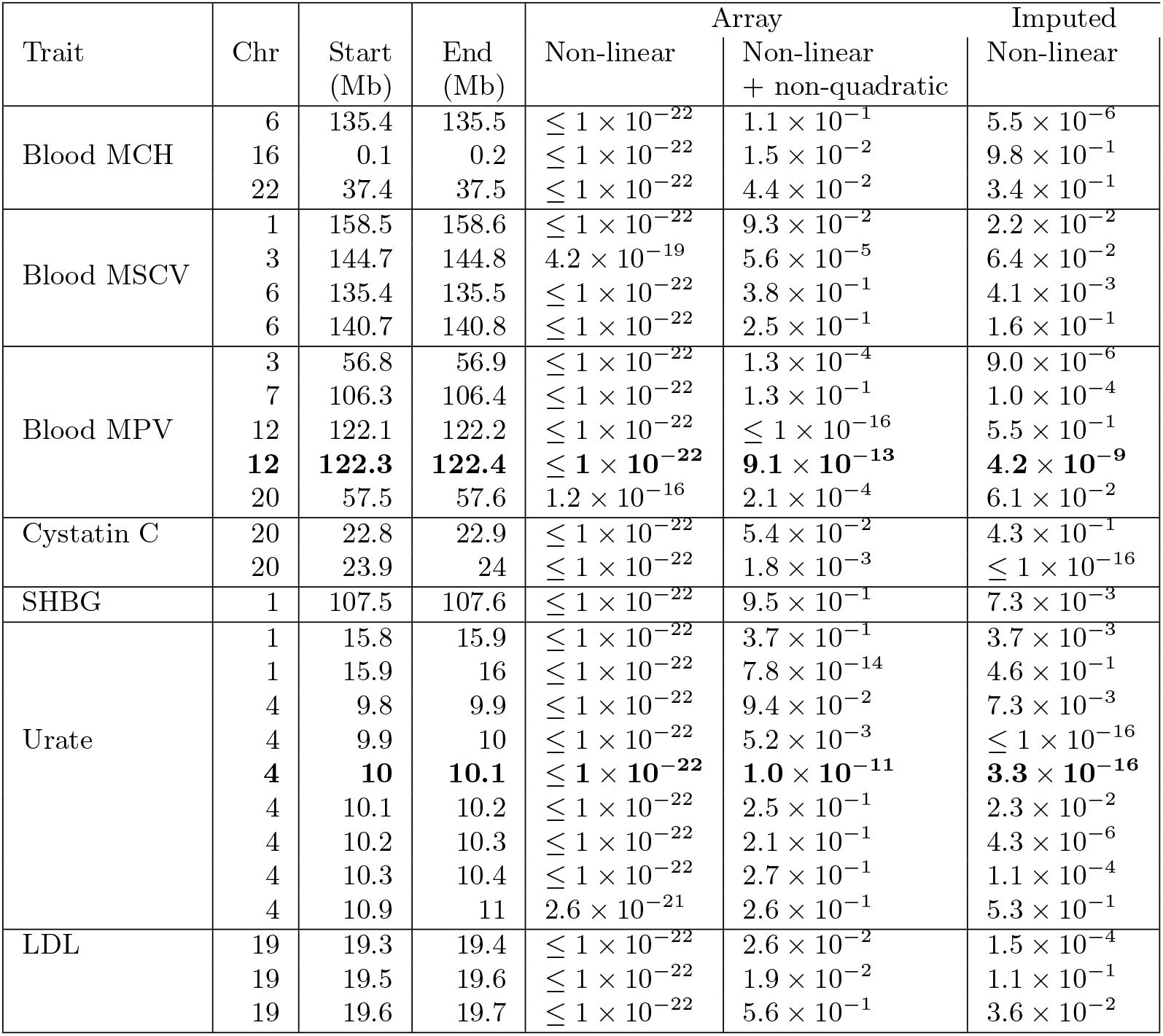
Loci with statistically significant non-linear effects (*p* <= 6 × 10^−8^ accounting for the number of sets and traits tested). We report loci with p-values that are significant after regressing out linear effects of SNPs in five windows centered around the tested window. We also report the p-value at these loci after removing the quadratic effect of the current window in addition to the linear effect. We also report the p-value after removing the linear effect when analyzing imputed genotypes in these windows. We highlight the loci which remain significant in array and imputed data and after regressing non-linear and non-quadratic effects.

The locus associated with blood MPV (12:122.3-122.4 Mb) overlaps the gene CFAP251 which contains multiple variants strongly associated with platelet volume [23–25] as well as multiple rare variants associated with male fertility [26]. The locus 4:9.9-10.1 Mb associated with serum urate levels overlaps the solute transporter gene SLC2A9 which harbors multiple variants associated with serum urate levels [27–31]. Variants in this gene have been found to have sex-specific effects on urate levels. Further, serum urate levels are positively correlated with body mass index (BMI) with suggestive evidence for modulation of the effect of variants in SLC2A9 on serum urate levels by BMI [32–34]. To investigate potential sex-specific differences in effects on serum urate levels, we separately analyzed this locus in men and women. We computed p-values at the hyperparameter value that attained the minimum p-value (*γ* = 0.1) in men and women using FastKAST. We obtain a p-value of 6.8 × 10^−4^ in men and 5.2 × 10^−7^ in women even though the number of men and women is comparable in our analyses (*N* = 132, 020 for men and *N* = 150, 496 for women). Overall, these results indicate that the loci that show evidence of non-linear effects harbor variants with significant marginal effects.

## Discussion

We have described FastKAST, a computationally efficient algorithm that is capable of testing for non-linear genetic effects in Biobank-scale data. FastKAST yields well-calibrated tests with little loss in power relative to an exact test. Applying FastKAST to 30 quantitative traits measured across ≈ 300K unrelated white British individuals in UKBB, we discovered 27 non-linear associations across seven traits of which two loci show strong evidence for higher-order non-linear effects.

We end with a discussion of the limitations of our approach and directions for future work. First, FastKAST is designed to analyze quantitative traits. FastKAST can be extended to binary traits in a straightforward manner using a generalized linear mixed model with a logistic link function [11–13]. Second, nonlinear interactions are represented in FastKAST using the class of shift-invariant kernels (which include the widely-used RBF kernel). FastKAST has the potential to be extended for a wider class of kernels using other randomized approximations, *e*.*g*., Nyström kernel approximation [35]. We leave a more detailed exploration of alternative kernels and approximations for future work. Third, our results are localized to fairly broad windows of size 100kb. While the application to windows of size 100 kb was motivated by computational considerations, we anticipate that we can apply FastKAST to smaller windows to further refine our current signals. Finally, we note that though we have shown strong evidence for the potential existence of higher order feature interactions, our results must be interpreted with caution. The interpretation of genetic interactions is conditioned on the number and quality of SNPs analyzed. It has been shown that apparent interactions in the data can be explained by linearity with missing SNPs [3, 22, 36]. We have attempted to address this issue by replicating the loci discovered on the imputed genotypes in UKBB. While the imputed genotypes contain the vast majority of SNPs with minor allele frequency > 0.1%, it is likely to be missing rare SNPs. The availability of whole-exome sequencing data in the UK Biobank (and other biobanks) will allow a more thorough investigation of these effects in protein-coding regions of the genome.

## Methods

Let ***y*** denote the vector of phenotypes measured on *N* individuals and ***Z*** denote the design matrix of genotypes over *M* SNPs that are desired to be tested. The goal is to test for association between the set of *M* SNPs and the phenotype.

We model the distribution of phenotypes, ***y***, as:

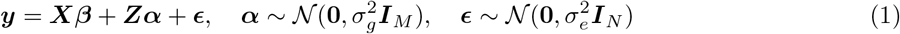

Here ***y*** ∈ ℝ^*N*^, ***X*** ∈ ℝ^*N* ×*P*^ denotes a matrix of covariates, ***Z*** ∈ ℝ^*N* ×*M*^ is the design matrix of standardized genotypes measured over *M* SNPs, and ***ϵ*** ℝ^*N*^ is the random vector of residual effects. ***β*** ∈ ℝ^*P*^ is the vector of fixed effect coefficients while ***α*** ∈ ℝ^*M*^ is the vector of random effect coefficients. 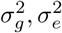 are the variance components associated with the genetic and residual effects. Integrating out the random effects, we have 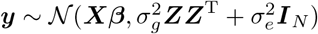.

The above model assumes that the genotype has a linear and additive effect on phenotype. To model non-linear effects, we transform the genotype using a non-linear function ***ϕ*** : ℝ^*M*^ → ℝ^*Q*^ leading to the following model:

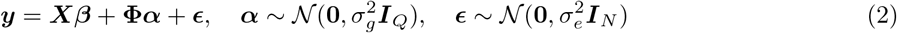

Here **Φ** is the design matrix obtained by applying ***ϕ***(***z***) to each individual genotype ***z. ϕ***(***z***) is assumed to lie in a Hilbert space endowed with a reproducing kernel function *k*(.,.) [37]. Equivalently, we can write this model as:

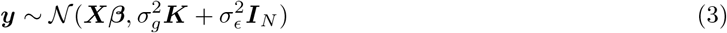

Here ***K*** is the *N* × *N* kernel matrix where *K*_*i,j*_ = *k*(***z***_*i*_, ***z***_*j*_), *i, j* ∈ {1, …, *N*}. For example, a common kernel is the radial basis function (RBF) kernel: 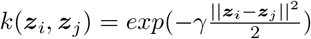.

### Hypothesis testing

Testing for a genetic contribution to the phenotype in Equation 3 involves testing the null hypothesis 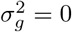. A commonly used approach to test the null hypothesis uses a score test [11]. Under the null hypothesis, the score statistic 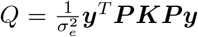 is asymptotically distributed as a weighted sum of 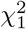 variables where the weights correspond to the eigenvalues of the matrix ***PKP*** and ***P*** = (***I*** − ***X***(***X***^*T*^ ***X***)^−1^***X***^*T*^) is the projection matrix. To compute the score statistic, an estimate of 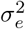, typically the restricted maximum likelihood (REML) estimate, is used. More recent works [38, 39] have characterized the sampling distribution of the score statistic in finite samples enabling the computation of exact p-values for the score test.

### Computation of p-values

A key challenge in computing p-values for the score statistic is the computation of all the eigenvalues of ***PKP*** which has time complexity 𝒪(*N*^3^). For weighted linear kernels, *i*.*e*., kernels of the form ***K*** = ***ZWZ***^T^ where ***W*** is a diagonal matrix with non-negative entries, this time complexity can be reduced to 𝒪(*NM*^2^). However, these approaches are not applicable to kernels that model general non-linear effects (like the RBF kernel) so that the computational complexity of testing for such effects scales as 𝒪(*N*^3^).

### Random Fourier Features

FastKAST relies on the observation that the kernel function can be approximated by mapping the input features to a randomized low-dimensional feature space [17]. For the class of shift-invariant kernel functions *k*(***z, z***′) = *f* (***z*** − ***z***′) (for some function *f*) that include the popular RBF kernel, the kernel function can be approximated by projecting each input ***z*** onto a random direction ***ω*** drawn from the Fourier transform of *k*. Specifically, we approximate 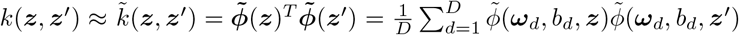. where *D* denotes the number of random features (which we term the *approximation dimension*), ***ω***_*d*_ ∈ ℝ^*M*^, *b*_*d*_ ∈ ℝ, 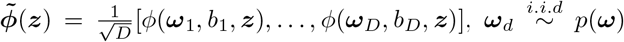 where *p*(***ω***) denotes the Fourier transform of *k*, 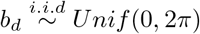, and 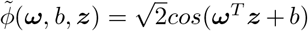. For example, in the RBF kernel with hyperparameter 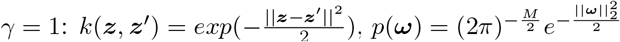, and in this case 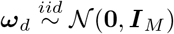.

Given the *N* × *D* approximate feature matrix 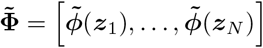, it follows that 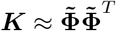. Prior work has shown that 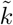 approximates *k* for a sufficiently large number of features *D* [17] (we empirically explore the approximation dimension *D* needed in our application). A key computational advantage of this approximation is that the approximate design matrix 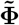 can be constructed in time linear in the sample size (𝒪(*NMD*)). We denote 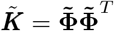 as the approximate kernel matrix.

### Hypothesis testing with random Fourier features

We use a score statistic to test the null hypothesis that 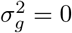 using the random Fourier feature approximation to the kernel. Let 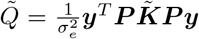 denote the approximate score statistic where ***P*** is the projection matrix. We show that, under the null hypothesis, the approximate score statistic is distributed as 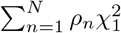 where *ρ*_*n*_ denotes the *n*^*th*^ eigenvalue of the matrix 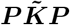 (Supplementary Note S1). We compute 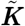 using random Fourier features while we estimate the noise variance 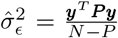. Computing p-values for the approximate score statistic 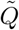 requires computing the eigenvalues of 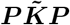 which can be computed from the SVD of 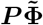 with time complexity 𝒪(*ND*^2^). Thus, the total time complexity of computing p-values using FastKAST is 𝒪(*NMD* + *ND*^2^).

### Computing p-values in the unknown hyperparameter setting

Applying FastKAST typically requires choosing a value for the kernel hyperparameter *γ*. First, we note that the hypothesis test remains calibrated for any choice of hyperparameter. However, the choice of hyperparameter can influence power. A naive approach to perform hypothesis tests while integrating over choices of the hyperparameter would involve selecting a set of hyperparameter values: {*γ*_1_, …, *γ*_*H*_} followed by computation of p-values *p*_*h*_ for each hyperparameter *h* (using the process described above). We then choose the minimum p-value: *p* = *min* {*p*_1_, …, *p*_*H*_} as the statistic. We then permute the trait and repeat the previous computation multiple times to approximate the sampling distribution of *p* (in the presence of covariates, we permute the residuals after fitting the covariates as fixed effects). We then compute the p-value of *p* using the sampling distribution approximated by the permutations. If the total number of permutation is *S*, then we need to compute *SH* score statistics for each set where *H* is the number of candidate hyperparameters. Given the precomputed eigenvalues of the approximate kernel matrix, the statistics for each permutation can be efficiently computed. However, the precision of the p-value is bounded by the 1*/S* requiring a large number of permutations to obtain accurate estimates of low p-values which makes the naive permutation approach computationally challenging. We therefore propose an approach similar to previous studies [18] to learn the null distribution of −log *p* values across sets using kernel density estimation (KDE) [40]. Under each set, we generate 10 different permutations, then we combine the corresponding statistics for all the sets to form the null distribution and learn it using kernel density estimation (KDE) with Gaussian kernel. After learning the distribution, we calculate the p-value from the estimated null distribution.

## Datasets

### Simulation dataset

We obtained a set of *N* = 337, 205 unrelated white British individuals measured at *M* = 593, 300 common SNPs genotyped on the UK Biobank Axiom array to use in simulations by extracting individuals that are > 3rd degree relatives and excluding individuals with putative sex chromosome aneuploidy.

### UKBB genotypes

For analysis of real traits, we restricted our analysis to SNPs that were presented in the UK Biobank Axiom array used to genotype the UK Biobank. SNPs with greater than 1% missingness and minor allele frequency smaller than 1% were removed. Moreover, SNPs that fail the Hardy-Weinberg test at significance threshold 10^−7^ were removed. We restricted our study to self-reported British white ancestry individuals which are > 3^*rd*^ degree relatives that is defined as pairs of individuals with kinship coefficient < 1/2^(9/2)^ [41]. Furthermore, we removed individuals who are outliers for genotype heterozygosity and/or missingness. Finally we obtained a set of *N* = 291, 273 individuals and *M* = 459, 792 SNPs to use in the real data analyses.

We also analyzed imputed genotypes across *N* = 291, 273 unrelated white British individuals. We removed SNPs with greater than 1% missingness, minor allele frequency smaller than 1%, SNPs that fail the Hardy-Weinberg test at significance threshold 10^−7^ as well as SNPs that lie within the MHC region (Chr6: 25–35 Mb) to obtain 4, 824, 392 SNPs.

### Covariates and phenotypes

We selected thirty quantitative traits in the UKBB which we processed using inverse rank-normalization. We included sex, age, and the top 20 genetic principal components (PCs) as covariates in our analysis for all phenotypes. We used PCs computed in the UKBB from a superset of 488, 295 individuals. Extra covariates were added for diastolic/systolic blood pressure (adjusted for cholesterol-lowering medication, blood pressure medication, insulin, hormone replacement therapy, and oral contraceptives) and waist-to-hip ratio (adjusted for BMI).

## Acknowledgments

This research was conducted using the UK Biobank Resource under application 33127. We thank the participants of UK Biobank for making this work possible.

## Code Availability

FastKAST can be found at https://github.com/sriramlab/FastKAST.

## References

[1] Snehit Prabhu and Itsik Pe’er. Ultrafast genome-wide scan for snp–snp interactions in common complex disease. Genome research, 22(11):2230–2240, 2012.

[2] Lars Wienbrandt, Jan Christian Kässens, Jorge González-Domínguez, Bertil Schmidt, David Ellinghaus, and Manfred Schimmler. Fpga-based acceleration of detecting statistical epistasis in gwas. Procedia Computer Science, 29:220–230, 2014.

[3] Gibran Hemani, Konstantin Shakhbazov, Harm-Jan Westra, Tonu Esko, Anjali K Henders, Allan F McRae, Jian Yang, Greg Gibson, Nicholas G Martin, Andres Metspalu, et al. Detection and replication of epistasis influencing transcription in humans. Nature, 508(7495):249–253, 2014.

[4] Wen-Hua Wei, Gibran Hemani, and Chris S Haley. Detecting epistasis in human complex traits. Nature Reviews Genetics, 15(11):722–733, 2014.

[5] Tobias L Lenz, Aaron J Deutsch, Buhm Han, Xinli Hu, Yukinori Okada, Stephen Eyre, Michael Knapp, Alexandra Zhernakova, Tom WJ Huizinga, Goncalo Abecasis, et al. Widespread non-additive and interaction effects within hla loci modulate the risk of autoimmune diseases. Nature genetics, 47(9):1085, 2015.

[6] Omer Weissbrod, Dan Geiger, and Saharon Rosset. Multikernel linear mixed models for complex phenotype prediction. Genome Research, 26(7):969–979, 2016.

[7] Lorin Crawford, Ping Zeng, Sayan Mukherjee, and Xiang Zhou. Detecting epistasis with the marginal epistasis test in genetic mapping studies of quantitative traits. PLoS genetics, 13(7):e1006869, 2017.

[8] Bingshan Li and Suzanne M Leal. Methods for detecting associations with rare variants for common diseases: application to analysis of sequence data. The American Journal of Human Genetics, 83(3):311–321, 2008.

[9] Bo Eskerod Madsen and Sharon R Browning. A groupwise association test for rare mutations using a weighted sum statistic. PLoS genetics, 5(2):e1000384, 2009.

[10] Benjamin M Neale, Manuel A Rivas, Benjamin F Voight, David Altshuler, Bernie Devlin, Marju Orho-Melander, Sekar Kathiresan, Shaun M Purcell, Kathryn Roeder, and Mark J Daly. Testing for an unusual distribution of rare variants. PLoS Genet, 7(3):e1001322, 2011.

[11] Michael C Wu, Seunggeun Lee, Tianxi Cai, Yun Li, Michael Boehnke, and Xihong Lin. Rare-variant association testing for sequencing data with the sequence kernel association test. The American Journal of Human Genetics, 89(1):82–93, 2011.

[12] Seunggeun Lee, Mary J Emond, Michael J Bamshad, Kathleen C Barnes, Mark J Rieder, Deborah A Nickerson, ESP Lung Project Team, David C Christiani, Mark M Wurfel, Xihong Lin, et al. Optimal unified approach for rare-variant association testing with application to small-sample case-control whole-exome sequencing studies. The American Journal of Human Genetics, 91(2):224–237, 2012.

[13] Iuliana Ionita-Laza, Seunggeun Lee, Vlad Makarov, Joseph D Buxbaum, and Xihong Lin. Sequence kernel association tests for the combined effect of rare and common variants. The American Journal of Human Genetics, 92(6):841–853, 2013.

[14] Clare Bycroft, Colin Freeman, Desislava Petkova, Gavin Band, Lloyd T Elliott, Kevin Sharp, Allan Motyer, Damjan Vukcevic, Olivier Delaneau, Jared O’Connell, et al. The uk biobank resource with deep phenotyping and genomic data. Nature, 562(7726):203–209, 2018.

[15] John Michael Gaziano, John Concato, Mary Brophy, Louis Fiore, Saiju Pyarajan, James Breeling, Stacey Whitbourne, Jennifer Deen, Colleen Shannon, Donald Humphries, et al. Million veteran program: a mega-biobank to study genetic influences on health and disease. Journal of clinical epidemiology, 70:214–223, 2016.

[16] Masahiro Kanai, Masato Akiyama, Atsushi Takahashi, Nana Matoba, Yukihide Momozawa, Masashi Ikeda, Nakao Iwata, Shiro Ikegawa, Makoto Hirata, Koichi Matsuda, et al. Genetic analysis of quantitative traits in the japanese population links cell types to complex human diseases. Nature genetics, 50(3):390–400, 2018.

[17] Ali Rahimi and Benjamin Recht. Random features for large-scale kernel machines. In J. C. Platt, D. Koller, Y. Singer, and S. T. Roweis, editors, Advances in Neural Information Processing Systems 20, pages 1177–1184. Curran Associates, Inc., 2008.

[18] Jennifer Listgarten, Christoph Lippert, Eun Yong Kang, Jing Xiang, Carl M Kadie, and David Heckerman. A powerful and efficient set test for genetic markers that handles confounders. Bioinformatics, 29(12):1526–1533, 2013.

[19] Peter M Visscher, Naomi R Wray, Qian Zhang, Pamela Sklar, Mark I McCarthy, Matthew A Brown, and Jian Yang. 10 years of gwas discovery: biology, function, and translation. The American Journal of Human Genetics, 101(1):5–22, 2017.

[20] Ali Pazokitoroudi, Alec M Chiu, Kathryn S Burch, Bogdan Pasaniuc, and Sriram Sankararaman. Quantifying the contribution of dominance deviation effects to complex trait variation in biobank-scale data. The American Journal of Human Genetics, 108(5):799–808, 2021.

[21] Valentin Hivert, Julia Sidorenko, Florian Rohart, Michael E Goddard, Jian Yang, Naomi R Wray, Loic Yengo, and Peter M Visscher. Estimation of non-additive genetic variance in human complex traits from a large sample of unrelated individuals. The American Journal of Human Genetics, 108(5):786–798, 2021.

[22] Frank Dudbridge and Olivia Fletcher. Gene-environment dependence creates spurious gene-environment interaction. The American Journal of Human Genetics, 95(3):301–307, 2014.

[23] Christa Meisinger, Holger Prokisch, Christian Gieger, Nicole Soranzo, Divya Mehta, Dieter Rosskopf, Peter Lichtner, Norman Klopp, Jonathan Stephens, Nicholas A Watkins, et al. A genome-wide association study identifies three loci associated with mean platelet volume. The American Journal of Human Genetics, 84(1):66–71, 2009.

[24] Nicole Soranzo, Tim D Spector, Massimo Mangino, Brigitte Kühnel, Augusto Rendon, Alexander Teumer, Christina Willenborg, Benjamin Wright, Li Chen, Mingyao Li, et al. A genome-wide metaanalysis identifies 22 loci associated with eight hematological parameters in the haemgen consortium. Nature genetics, 41(11):1182–1190, 2009.

[25] William J Astle, Heather Elding, Tao Jiang, Dave Allen, Dace Ruklisa, Alice L Mann, Daniel Mead, Heleen Bouman, Fernando Riveros-Mckay, Myrto A Kostadima, et al. The allelic landscape of human blood cell trait variation and links to common complex disease. Cell, 167(5):1415–1429, 2016.

[26] Weiyu Li, Xiaojin He, Shenmin Yang, Chunyu Liu, Huan Wu, Wangjie Liu, Mingrong Lv, Dongdong Tang, Jing Tan, Shuyan Tang, et al. Biallelic mutations of cfap251 cause sperm flagellar defects and human male infertility. Journal of human genetics, 64(1):49–54, 2019.

[27] Angela Döring, Christian Gieger, Divya Mehta, Henning Gohlke, Holger Prokisch, Stefan Coassin, Guido Fischer, Kathleen Henke, Norman Klopp, Florian Kronenberg, et al. Slc2a9 influences uric acid concentrations with pronounced sex-specific effects. Nature genetics, 40(4):430–436, 2008.

[28] Veronique Vitart, Igor Rudan, Caroline Hayward, Nicola K Gray, James Floyd, Colin NA Palmer, Sara A Knott, Ivana Kolcic, Ozren Polasek, Juergen Graessler, et al. Slc2a9 is a newly identified urate transporter influencing serum urate concentration, urate excretion and gout. Nature genetics, 40(4):437–442, 2008.

[29] Anna Köttgen, Eva Albrecht, Alexander Teumer, Veronique Vitart, Jan Krumsiek, Claudia Hundertmark, Giorgio Pistis, Daniela Ruggiero, Conall M O’Seaghdha, Toomas Haller, et al. Genome-wide association analyses identify 18 new loci associated with serum urate concentrations. Nature genetics, 45(2):145–154, 2013.

[30] Nasa Sinnott-Armstrong, Yosuke Tanigawa, David Amar, Nina Mars, Christian Benner, Matthew Aguirre, Guhan Ram Venkataraman, Michael Wainberg, Hanna M Ollila, Tuomo Kiiskinen, et al. Genetics of 35 blood and urine biomarkers in the uk biobank. Nature genetics, 53(2):185–194, 2021.

[31] Yoichiro Kamatani, Koichi Matsuda, Yukinori Okada, Michiaki Kubo, Naoya Hosono, Yataro Daigo, Yusuke Nakamura, and Naoyuki Kamatani. Genome-wide association study of hematological and biochemical traits in a japanese population. Nature genetics, 42(3):210–215, 2010.

[32] Anita Brandstätter, Stefan Kiechl, Barbara Kollerits, Steven C Hunt, Iris M Heid, Stefan Coassin, Johann Willeit, Ted D Adams, Thomas Illig, Paul N Hopkins, et al. Sex-specific association of the putative fructose transporter slc2a9 variants with uric acid levels is modified by bmi. Diabetes care, 31(8):1662–1667, 2008.

[33] Wei-Dong Li, Hongxiao Jiao, Kai Wang, Clarence K Zhang, Joseph T Glessner, Struan FA Grant, Hongyu Zhao, Hakon Hakonarson, and R Arlen Price. A genome wide association study of plasma uric acid levels in obese cases and never-overweight controls. Obesity, 21(9):E490–E494, 2013.

[34] Jennifer E Huffman, Eva Albrecht, Alexander Teumer, Massimo Mangino, Karen Kapur, Toby Johnson, Zoltán Kutalik, Nicola Pirastu, Giorgio Pistis, Lorna M Lopez, et al. Modulation of genetic associations with serum urate levels by body-mass-index in humans. PloS one, 10(3):e0119752, 2015.

[35] Petros Drineas, Michael W Mahoney, and Nello Cristianini. On the nyström method for approximating a gram matrix for improved kernel-based learning. journal of machine learning research, 6(12), 2005.

[36] Andrew R Wood, Marcus A Tuke, Mike A Nalls, Dena G Hernandez, Stefania Bandinelli, Andrew B Singleton, David Melzer, Luigi Ferrucci, Timothy M Frayling, and Michael N Weedon. Another explanation for apparent epistasis. Nature, 514(7520):E3–E5, 2014.

[37] John Shawe-Taylor and Nello Cristianini. Kernel Methods for Pattern Analysis. Cambridge University Press, 2004.

[38] Han Chen, Thomas Lumley, Jennifer Brody, Nancy L Heard-Costa, Caroline S Fox, L Adrienne Cupples, and Josée Dupuis. Sequence kernel association test for survival traits. Genetic epidemiology, 38(3):191–197, 2014.

[39] Regev Schweiger, Omer Weissbrod, Elior Rahmani, Martina Müller-Nurasyid, Sonja Kunze, Christian Gieger, Melanie Waldenberger, Saharon Rosset, and Eran Halperin. Rl-skat: an exact and efficient score test for heritability and set tests. Genetics, 207(4):1275–1283, 2017.

[40] David W Scott. Multivariate density estimation: theory, practice, and visualization. John Wiley & Sons, 2015.

[41] Clare Bycroft, Colin Freeman, Desislava Petkova, Gavin Band, Lloyd T Elliott, Kevin Sharp, Allan Motyer, Damjan Vukcevic, Olivier Delaneau, Jared O’Connell, et al. The uk biobank resource with deep phenotyping and genomic data. Nature, 562(7726):203–209, 2018.

