## Supplementary information for "Fast Kernel-based Association Testing of non-linear genetic effects for Biobank-scale data"

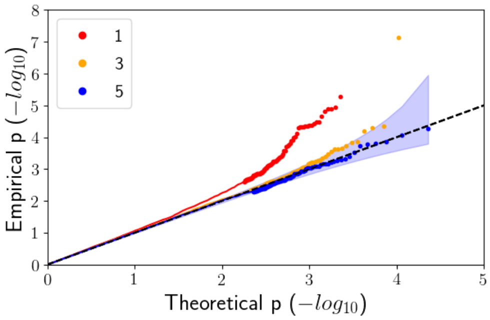


Figure S1: **Accounting for linear effects in the application of FastKAST**. We regressed linear effects in windows surrounding the target window (superwindow) that is being tested. We varied the number of windows comprising the superwindow (equal to the target window, 3 windows, and 5 windows surrounding the target window) to find that a superwindow of size 5 leads to calibrated p-values. The phenotypes were simulated under the non-infinitesimal genetic architecture with causal ratio = 0.001 and RARE distribution of causal variants.

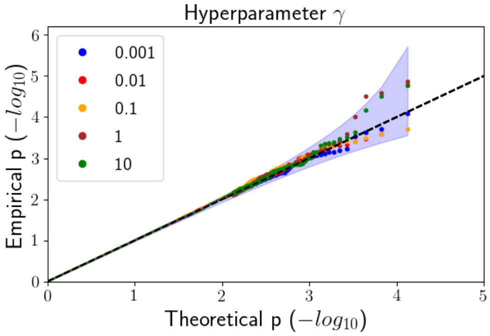


Figure S2: **Calibration of FastKAST under the choice of specific values for kernel hyperparameters**. We assessed the calibration of FastKAST with different kernel hyperparameters under simulations with linear effects but no nonlinear effect. We varied the kernel hyperparameter $\gamma$ in simulations with heritability $h^2 = 0.5$, causal variant ratio = 0.001, and the MAF range of causal variants ranging from 0 to 0.5 (ALL). We fit FastKAST, in turn, with RBF kernel hyperparameter: ($\gamma \in \{0.001, 0.01, 0.1, 1, 10\}$).

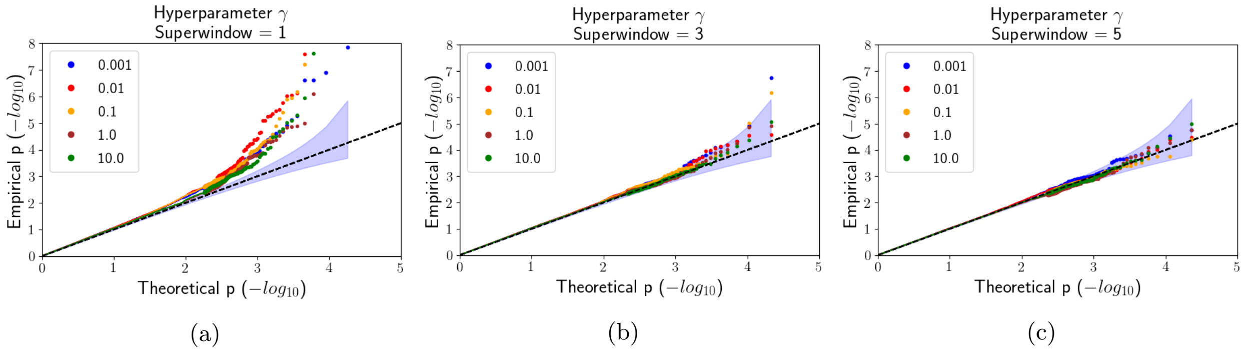


Figure S3: **Accounting for linear effects in the application of FastKAST**. We regressed linear effects in windows surrounding the target window that is being tested. We varied the number of windows surrounding the target window to find that regressing linear effects in the five windows surrounding the target window leads to calibrated p-values. The architecture that we simulate under here is RARE with causal ratio = 0.001.

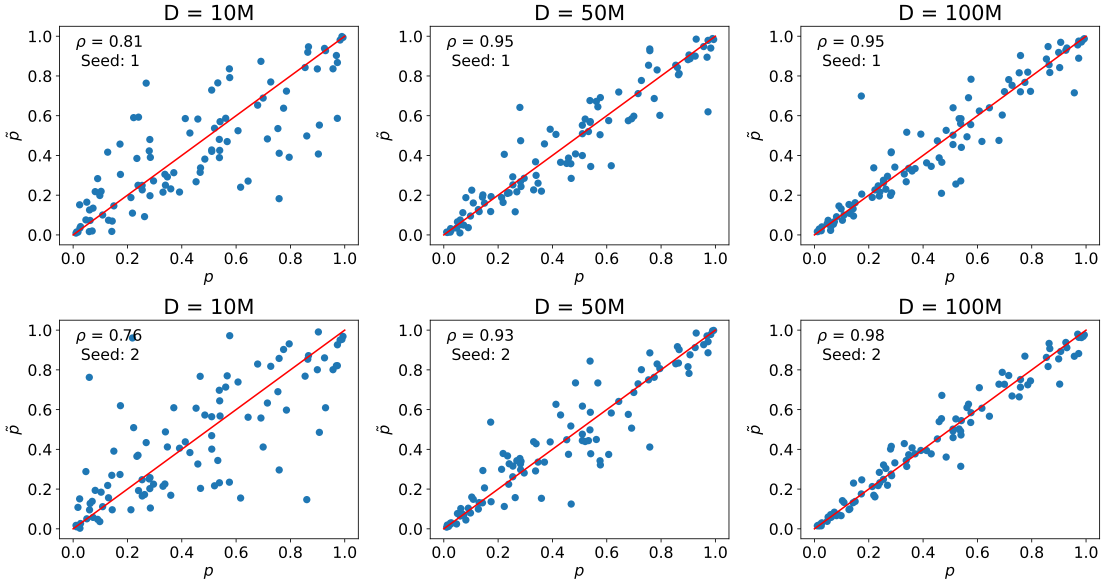


Figure S4: **Correlation between p value from approximating kernel (for two different random seeds) and true kernel using simulated phenotype**. $p$ represents the p-value computed using the exact kernel; $\tilde{p}$ represents the p-value computed by FastKAST as a function of the approximation dimension.

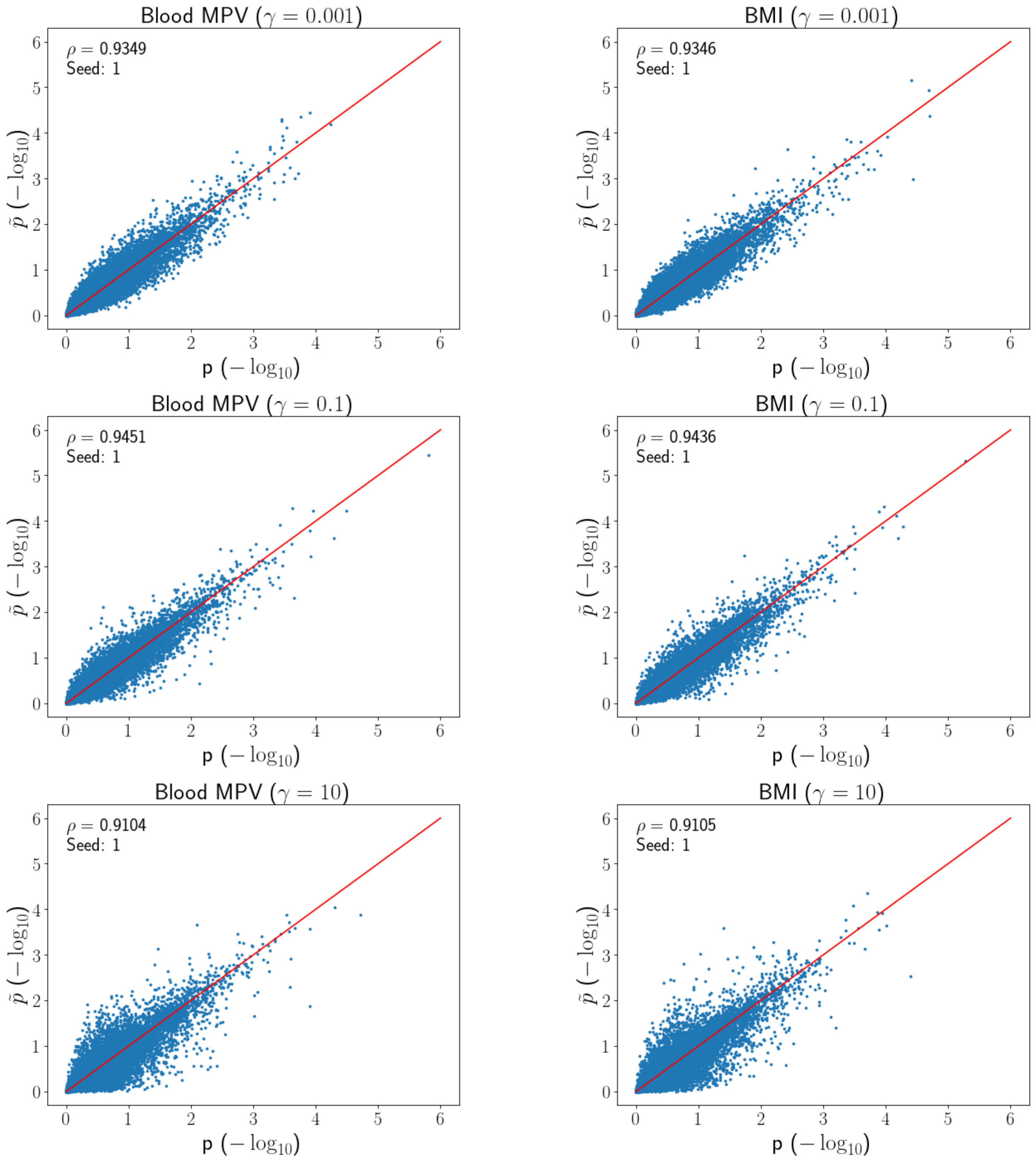


Figure S5: **Correlation between p value from approximating kernel (for two different random seeds) and true kernel using real trait phenotype under various kernel hyperparameter values ($\gamma$).** $p$ represents the p-value computed using the exact kernel; $\tilde{p}$ represents the p-value computed by FastKAST as a function of the approximation dimension.

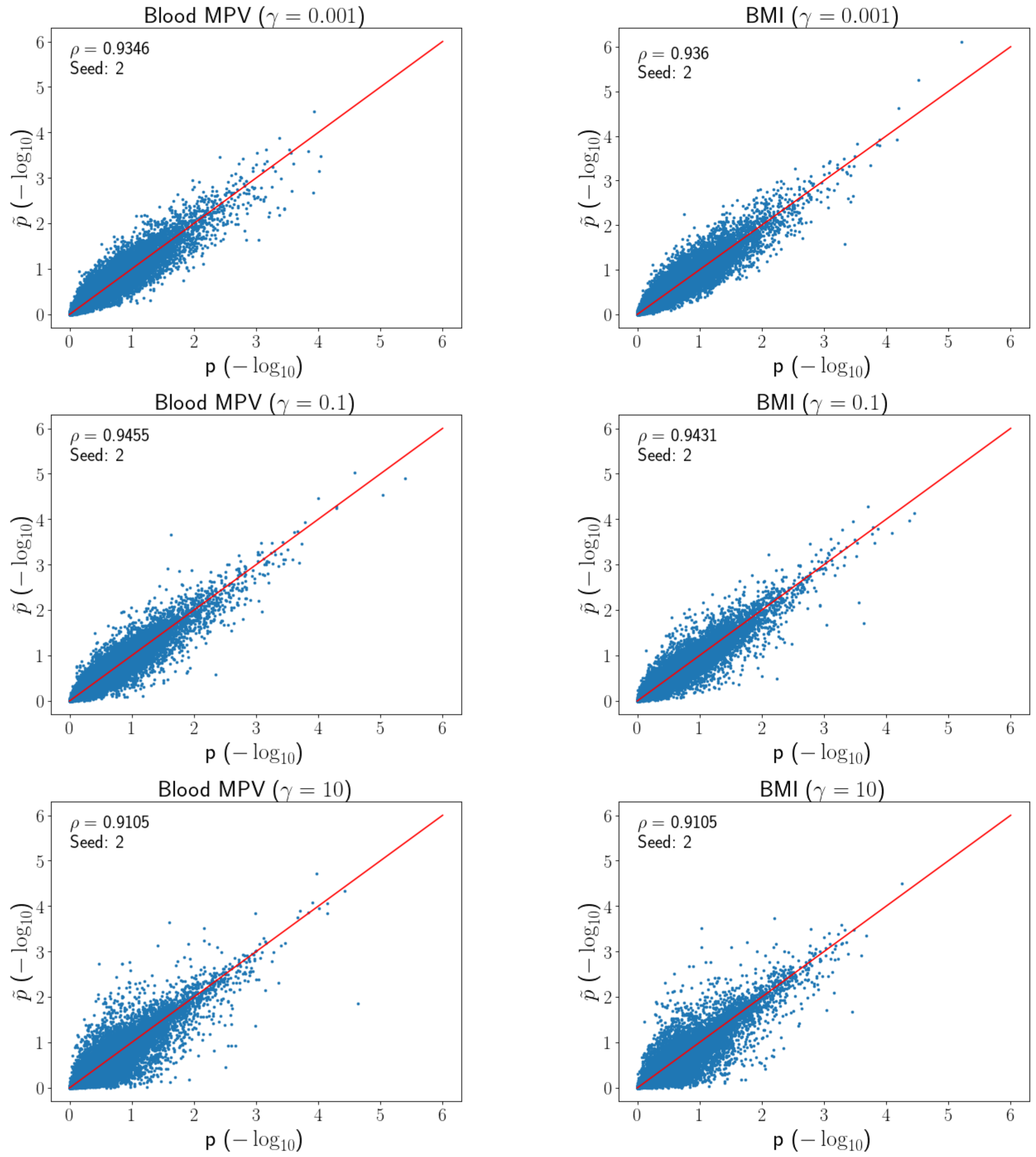


Figure S6: **Correlation between p value from approximating kernel (for two different random seeds) and true kernel using real trait phenotype under various kernel hyperparameter values ($\gamma$) (seed 2)**. $p$ represents the p-value computed using the exact kernel; $\tilde{p}$ represents the p-value computed by FastKAST as a function of the approximation dimension.

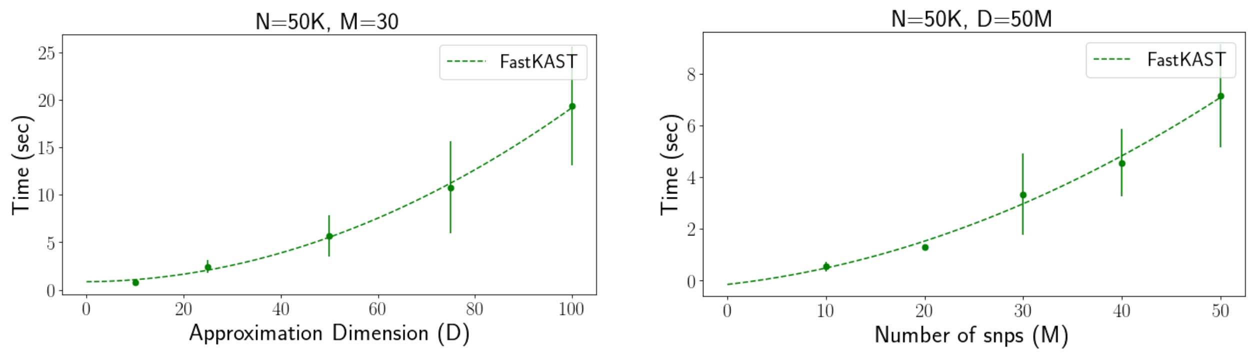


(a) (b)

Figure S7: **Computational efficiency of FastKAST**. (a) shows the runtime of FastKAST when increasing the approximation dimension $D$. (b) shows the runtime of FastKAST when increasing the number of SNPs $M$ within the window. The figure only demonstrates the computational time of the singular value decomposition (SVD) which is the bottleneck of FastKAST.

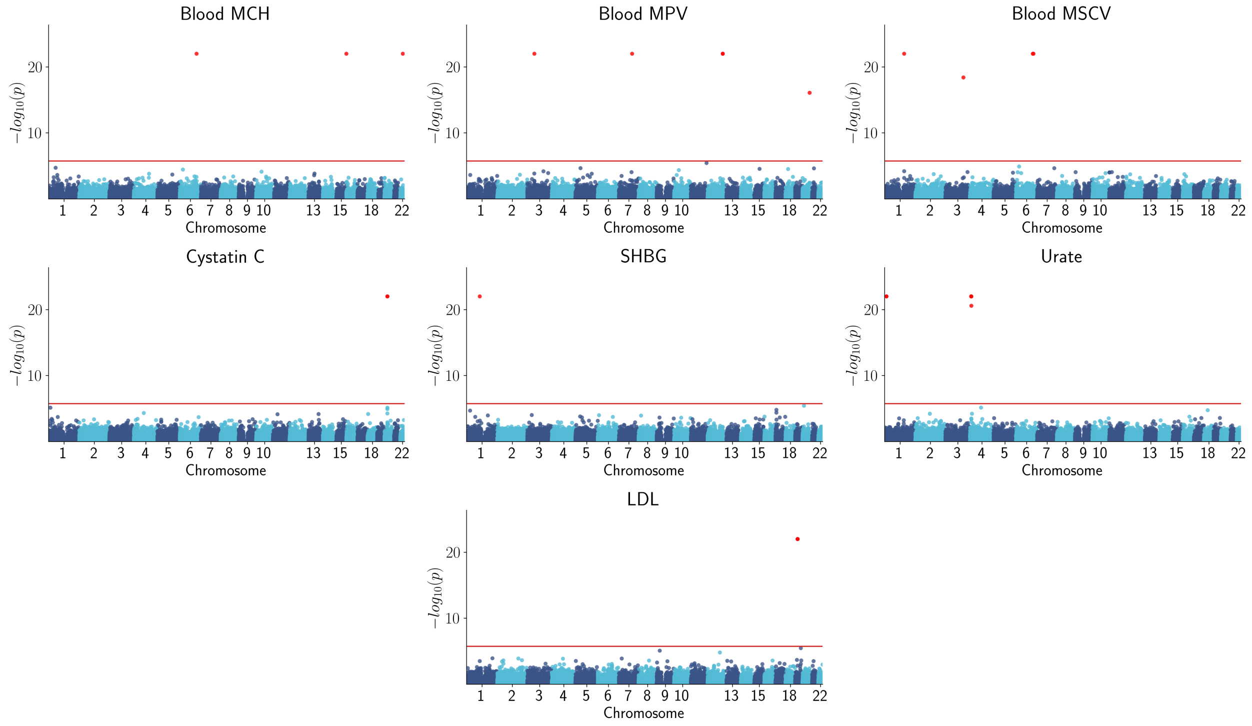


Figure S8: **Manhattan plot for test of non-linear effects**

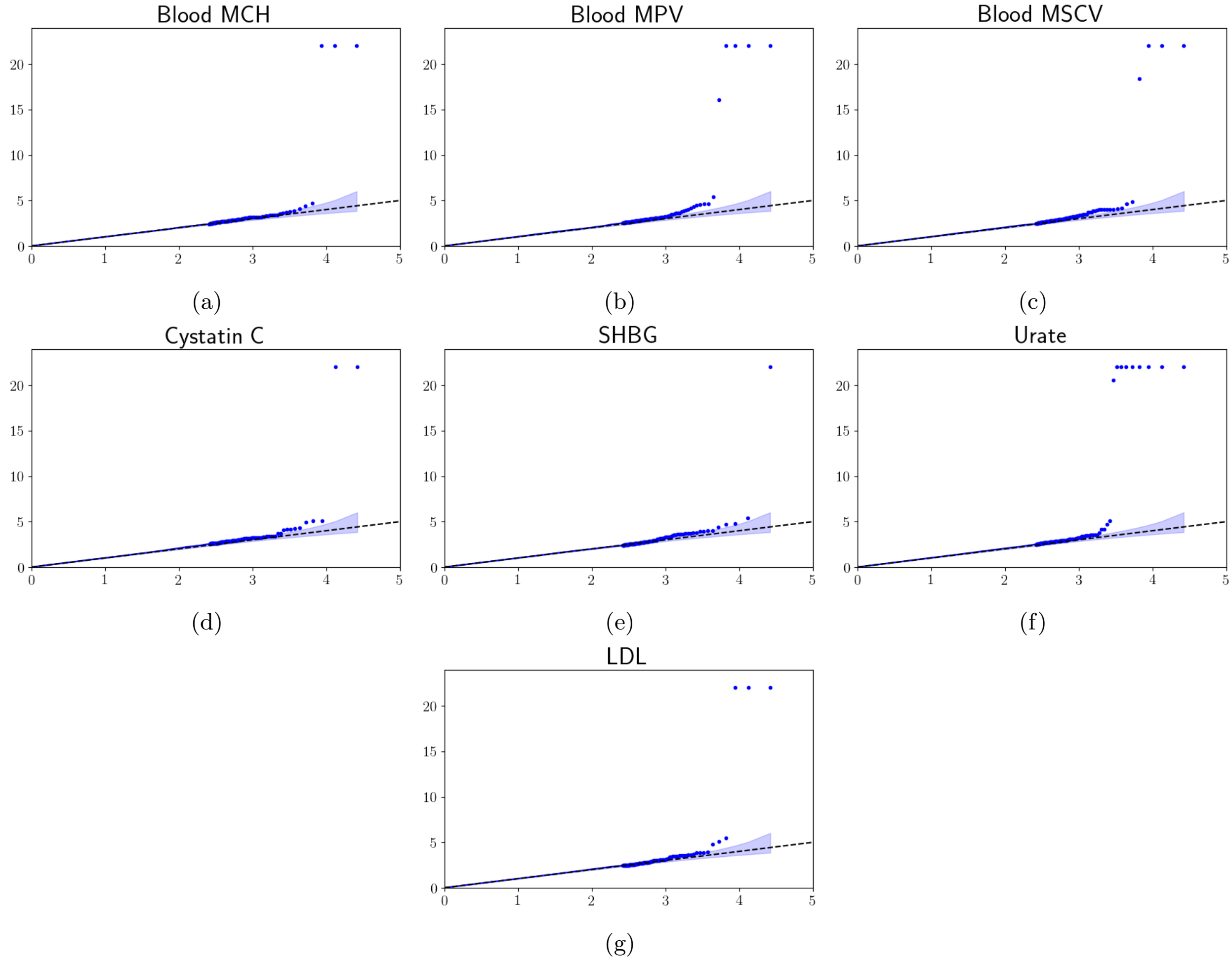


Figure S9: **Tests for non-linear effects in traits in the UK Biobank**. We used FastKAST to test for non-linear effects within 100 kb windows in 30 quantitative traits measured in unrelated white British individuals in the UK Biobank ($N = 291,273$, $M = 459,792$ array SNPs). In all of these analyses, we regressed out linear effects within five windows surrounding the tested window. We plot seven traits for which at least one window passed the Bonferroni significance threshold (a total of $27,899$ sets located in contiguous regions).

| Superwindow | $10^{-1}$ | $10^{-2}$ | $10^{-3}$ | $10^{-4}$ | $10^{-5}$ | $\lambda_{gc}$ |
|---|---|---|---|---|---|---|
| 1 | 0.113 | 0.015 | 0.003 | 1.450e-03 | 4.461e-04 | 1.093 |
| 3 | 0.103 | 0.011 | 0.001 | 2.356e-04 | 9.426e-05 | 1.031 |
| 5 | 0.101 | 0.010 | 0.001 | 8.718e-05 | 0.000e+00 | 1.021 |

Table S1: **False positive rate for varying p-value thresholds and genomic inflation factor**. We report the false positive rate and genomic inflation factor at different p-value thresholds (when the hyperparameter is adaptively chosen) as we vary the number of neighboring windows used to regress out linear effects (superwindow).

| Superwindow | $\gamma$ | $10^{-1}$ | $10^{-2}$ | $10^{-3}$ | $10^{-4}$ | $10^{-5}$ | $\lambda_{gc}$ |
|---|---|---|---|---|---|---|---|
| 1 | 0.001 | 0.116 | 0.015 | 2.737e-03 | 8.939e-04 | 3.352e-04 | 1.111 |
| 1 | 0.010 | 0.116 | 0.015 | 3.516e-03 | 1.507e-03 | 8.372e-04 | 1.117 |
| 1 | 0.100 | 0.112 | 0.016 | 2.621e-03 | 1.004e-03 | 5.020e-04 | 1.097 |
| 1 | 1.000 | 0.110 | 0.011 | 2.511e-03 | 7.812e-04 | 1.674e-04 | 1.081 |
| 1 | 10.000 | 0.111 | 0.012 | 2.400e-03 | 7.254e-04 | 3.906e-04 | 1.084 |
| 3 | 0.001 | 0.107 | 0.011 | 1.227e-03 | 2.832e-04 | 4.721e-05 | 1.042 |
| 3 | 0.010 | 0.108 | 0.011 | 1.038e-03 | 2.830e-04 | 0.000e+00 | 1.065 |
| 3 | 0.100 | 0.104 | 0.011 | 1.367e-03 | 1.414e-04 | 9.428e-05 | 1.066 |
| 3 | 1.000 | 0.105 | 0.011 | 8.488e-04 | 1.886e-04 | 0.000e+00 | 1.028 |
| 3 | 10.000 | 0.106 | 0.011 | 1.037e-03 | 1.415e-04 | 4.716e-05 | 1.046 |
| 5 | 0.001 | 0.104 | 0.011 | 1.048e-03 | 1.310e-04 | 0.000e+00 | 1.041 |
| 5 | 0.010 | 0.106 | 0.011 | 7.851e-04 | 1.309e-04 | 0.000e+00 | 1.043 |
| 5 | 0.100 | 0.102 | 0.010 | 8.283e-04 | 4.360e-05 | 0.000e+00 | 1.032 |
| 5 | 1.000 | 0.102 | 0.010 | 7.850e-04 | 1.744e-04 | 0.000e+00 | 1.023 |
| 5 | 10.000 | 0.102 | 0.010 | 8.723e-04 | 1.308e-04 | 0.000e+00 | 1.024 |

Table S2: **False positive rate for varying p-value thresholds and genomic inflation factor**. We report the false positive rate and genomic inflation factor at different p-value thresholds as we vary the hyperparameter and the number of windows used to regress out linear effects (superwindow).

| Trait | Chr | Start (Mb) | End (Mb) | $\gamma$ | Number of PCs | | | |
|---|---|---|---|---|---|---|---|---|
| | | | | | 5 | 10 | 20 | 40 |
| Blood MCH | 6 | 135.4 | 135.5 | 0.001 | $\leq 1 \times 10^{-14}$ | $6.8 \times 10^{-15}$ | $\leq 1 \times 10^{-14}$ | $\leq 1 \times 10^{-14}$ |
| Blood MCH | 16 | 0.1 | 0.2 | 0.01 | $2.4 \times 10^{-13}$ | $3.2 \times 10^{-13}$ | $2.9 \times 10^{-13}$ | $2.9 \times 10^{-13}$ |
| Blood MCH | 22 | 37.4 | 37.5 | 0.01 | $2.2 \times 10^{-8}$ | $1.9 \times 10^{-8}$ | $1.2 \times 10^{-8}$ | $1.2 \times 10^{-8}$ |
| Blood MSCV | 1 | 158.5 | 158.6 | 1.0 | $1.4 \times 10^{-11}$ | $1.5 \times 10^{-11}$ | $1.5 \times 10^{-11}$ | $1.5 \times 10^{-11}$ |
| Blood MSCV | 3 | 144.7 | 144.8 | 10.0 | $1.5 \times 10^{-6}$ | $1.5 \times 10^{-6}$ | $1.4 \times 10^{-6}$ | $1.4 \times 10^{-6}$ |
| Blood MSCV | 6 | 135.4 | 135.5 | 0.001 | $1.4 \times 10^{-6}$ | $1.3 \times 10^{-6}$ | $1.2 \times 10^{-6}$ | $1.2 \times 10^{-6}$ |
| Blood MSCV | 6 | 140.7 | 140.8 | 0.01 | $2.9 \times 10^{-8}$ | $2.4 \times 10^{-8}$ | $3.2 \times 10^{-8}$ | $2.5 \times 10^{-8}$ |
| Blood MPV | 3 | 56.8 | 56.9 | 0.01 | $2.3 \times 10^{-15}$ | $\leq 1 \times 10^{-14}$ | $\leq 1 \times 10^{-14}$ | $\leq 1 \times 10^{-14}$ |
| Blood MPV | 7 | 106.3 | 106.4 | 0.01 | $1.5 \times 10^{-8}$ | $1.3 \times 10^{-8}$ | $1.4 \times 10^{-8}$ | $1.4 \times 10^{-8}$ |
| Blood MPV | 12 | 122.1 | 122.2 | 10.0 | $2.1 \times 10^{-9}$ | $2.1 \times 10^{-9}$ | $2.2 \times 10^{-9}$ | $2.3 \times 10^{-9}$ |
| Blood MPV | 12 | 122.3 | 122.4 | 0.1 | $1.6 \times 10^{-11}$ | $1.6 \times 10^{-11}$ | $1.6 \times 10^{-11}$ | $1.7 \times 10^{-11}$ |
| Blood MPV | 20 | 57.5 | 57.6 | 0.01 | $3.2 \times 10^{-7}$ | $3.5 \times 10^{-7}$ | $3.6 \times 10^{-7}$ | $3.2 \times 10^{-7}$ |
| Cystatin C | 20 | 22.8 | 22.8 | 0.1 | $3.1 \times 10^{-7}$ | $3.4 \times 10^{-7}$ | $2.9 \times 10^{-7}$ | $2.9 \times 10^{-7}$ |
| Cystatin C | 20 | 23.9 | 24 | 0.01 | $6.1 \times 10^{-12}$ | $8.6 \times 10^{-12}$ | $1.2 \times 10^{-11}$ | $1.6 \times 10^{-11}$ |
| SHBG | 1 | 107.5 | 107.6 | 0.1 | $7.2 \times 10^{-10}$ | $7.2 \times 10^{-10}$ | $8.6 \times 10^{-10}$ | $8.7 \times 10^{-10}$ |
| Urate | 1 | 15.8 | 15.9 | 0.01 | $9.8 \times 10^{-13}$ | $8.8 \times 10^{-13}$ | $1.4 \times 10^{-12}$ | $1.3 \times 10^{-12}$ |
| Urate | 1 | 15.9 | 16 | 1.0 | $1.6 \times 10^{-12}$ | $1.5 \times 10^{-12}$ | $2.6 \times 10^{-12}$ | $2.9 \times 10^{-12}$ |
| Urate | 4 | 9.8 | 9.9 | 0.01 | $7.2 \times 10^{-10}$ | $7.6 \times 10^{-10}$ | $7.2 \times 10^{-10}$ | $7.4 \times 10^{-10}$ |
| Urate | 4 | 9.9 | 10 | 0.001 | $\leq 1 \times 10^{-14}$ | $\leq 1 \times 10^{-14}$ | $\leq 1 \times 10^{-14}$ | $\leq 1 \times 10^{-14}$ |
| Urate | 4 | 10 | 10.1 | 0.001 | $\leq 1 \times 10^{-14}$ | $\leq 1 \times 10^{-14}$ | $\leq 1 \times 10^{-14}$ | $\leq 1 \times 10^{-14}$ |
| Urate | 4 | 10.1 | 10.2 | 0.001 | $\leq 1 \times 10^{-14}$ | $\leq 1 \times 10^{-14}$ | $\leq 1 \times 10^{-14}$ | $\leq 1 \times 10^{-14}$ |
| Urate | 4 | 10.2 | 10.3 | 0.001 | $\leq 1 \times 10^{-14}$ | $\leq 1 \times 10^{-14}$ | $\leq 1 \times 10^{-14}$ | $\leq 1 \times 10^{-14}$ |
| Urate | 4 | 10.3 | 10.4 | 0.001 | $\leq 1 \times 10^{-14}$ | $\leq 1 \times 10^{-14}$ | $\leq 1 \times 10^{-14}$ | $\leq 1 \times 10^{-14}$ |
| Urate | 4 | 10.9 | 11 | 0.01 | $9.0 \times 10^{-8}$ | $9.3 \times 10^{-8}$ | $1.1 \times 10^{-7}$ | $1.0 \times 10^{-7}$ |
| LDL | 19 | 19.3 | 19.4 | 0.01 | $7.8 \times 10^{-10}$ | $7.8 \times 10^{-10}$ | $6.8 \times 10^{-10}$ | $7.1 \times 10^{-10}$ |
| LDL | 19 | 19.5 | 19.6 | 0.01 | $4.0 \times 10^{-11}$ | $3.9 \times 10^{-11}$ | $3.8 \times 10^{-11}$ | $4.2 \times 10^{-11}$ |
| LDL | 19 | 19.6 | 19.7 | 0.01 | $\leq 1 \times 10^{-14}$ | $\leq 1 \times 10^{-14}$ | $\leq 1 \times 10^{-14}$ | $\leq 1 \times 10^{-14}$ |

Table S3: **Robustness with respect to population structure (Number of principal components)**. We test the significant loci with varying number of principal components. We report the p-value for the hyperparameter that attains the minimum p-value in our initial analysis for each locus.

### S1 Proof of the sampling distribution of the approximate score statistic

**Theorem 1.** *Assuming that* $\boldsymbol{y} \sim \mathcal{N}(\boldsymbol{X\beta}, \sigma_g^2\boldsymbol{K} + \sigma_\epsilon^2\boldsymbol{I})$*, the approximate score statistic* $Q = \frac{1}{\sigma_\epsilon^2}\boldsymbol{y}^T\boldsymbol{P\tilde{K}Py}$ *is distributed as a weighted sum of* $\chi_1^2$ *variables under the null hypothesis* ($H_0$)*:*

$$\frac{1}{\sigma_\epsilon^2}\boldsymbol{y}^T\boldsymbol{P\tilde{K}Py} \sim \sum_{n=1}^{N}\rho_n\chi_1^2$$

*where* $\rho_n$ *denotes the* $n^{th}$ *eigenvalue of the matrix* $\boldsymbol{P\tilde{K}P}$ *and* $\boldsymbol{P} = (\boldsymbol{I} - \boldsymbol{X}(\boldsymbol{X}^T\boldsymbol{X})^{-1}\boldsymbol{X}^T)$ *denotes the projection matrix.*

*Proof.* Under the null hypothesis $\sigma_g^2 = 0$, we have $\boldsymbol{y} \sim \mathcal{N}(\boldsymbol{X\beta}, \sigma_\epsilon^2\boldsymbol{I})$, therefore $\boldsymbol{Py} \sim \mathcal{N}(0, \sigma_\epsilon^2\boldsymbol{P})$. Since $\boldsymbol{\tilde{K}}$ is positive semi-definite, it has a unique square root $\boldsymbol{\tilde{K}}^{\frac{1}{2}}$. Thus $\boldsymbol{\tilde{K}}^{\frac{1}{2}}\boldsymbol{Py} \sim \mathcal{N}(0, \sigma_\epsilon^2\boldsymbol{\tilde{K}}^{\frac{1}{2}}\boldsymbol{P}(\boldsymbol{\tilde{K}}^{\frac{1}{2}})^T)$. Since $\boldsymbol{P}$ is symmetric and idempotent, we have $\boldsymbol{P} = \boldsymbol{P}^2 = \boldsymbol{PP}^{\mathrm{T}}$ so that:

$$\begin{aligned}\boldsymbol{\tilde{K}}^{\frac{1}{2}}\boldsymbol{P}(\boldsymbol{\tilde{K}}^{\frac{1}{2}})^T &= \boldsymbol{\tilde{K}}^{\frac{1}{2}}\boldsymbol{PP}^T(\boldsymbol{\tilde{K}}^{\frac{1}{2}})^T \\ &= \left(\boldsymbol{\tilde{K}}^{\frac{1}{2}}\boldsymbol{P}\right)\left(\boldsymbol{\tilde{K}}^{\frac{1}{2}}\boldsymbol{P}\right)^T\end{aligned}$$

Thus, $\boldsymbol{\tilde{K}}^{\frac{1}{2}}\boldsymbol{P}(\boldsymbol{\tilde{K}}^{\frac{1}{2}})^T \succeq 0$ so that $\boldsymbol{\tilde{K}}^{\frac{1}{2}}\boldsymbol{P}(\boldsymbol{\tilde{K}}^{\frac{1}{2}})^T = \boldsymbol{U\Sigma U}^T$ where $\boldsymbol{\Sigma} = diag(\rho_1, \ldots, \rho_N)$ and $\boldsymbol{U}^T\boldsymbol{U} = \boldsymbol{I}$. Setting $\boldsymbol{B} = \boldsymbol{U}^T\boldsymbol{\tilde{K}}^{\frac{1}{2}}\boldsymbol{P}$, we have $\boldsymbol{By} \sim \mathcal{N}(0, \sigma_\epsilon^2\boldsymbol{\Sigma})$. We then have:

$$\begin{aligned}Q &\equiv \frac{1}{\sigma_\epsilon^2}\boldsymbol{y}^T\boldsymbol{P\tilde{K}Py} \\ &= \frac{1}{\sigma_\epsilon^2}(\boldsymbol{U\tilde{K}}^{\frac{1}{2}}\boldsymbol{Py})^T(\boldsymbol{U\tilde{K}}^{\frac{1}{2}}\boldsymbol{Py}) = \frac{1}{\sigma_\epsilon^2}(\boldsymbol{By})^T\boldsymbol{By} \sim \sum_{n=1}^{N}\rho_n\chi_1^2\end{aligned}$$

The eigenvalues of $\boldsymbol{K}^{\frac{1}{2}}\boldsymbol{P}(\boldsymbol{K}^{\frac{1}{2}})^T$ are equal to the eigenvalues of $\boldsymbol{U}^T\boldsymbol{K}^{\frac{1}{2}}\boldsymbol{PP}(\boldsymbol{K}^{\frac{1}{2}})^T\boldsymbol{U} = \boldsymbol{BB}^T$. Using the fact that for every matrix $\boldsymbol{B}$, $\boldsymbol{B}^T\boldsymbol{B}$ and $\boldsymbol{BB}^T$ have the same nonzero eigenvalues, we can equivalently calculate $\rho_n$ using the $n^{th}$ eigenvalue of the matrix $\boldsymbol{P\tilde{K}P}$. □
